# Karyotype evolution of angel insects (Zoraptera)

**DOI:** 10.64898/2026.06.04.730103

**Authors:** Marek Jankásek, Ivona Kočárková, Petr Kočárek, František Št’áhlavský

**Affiliations:** Department of Zoology, Charles University, Viničná 7, 128 00 Prague, Czech Republic; Department of Biology and Ecology, University of Ostrava, 30. dubna 22, 701 03 Ostrava, Czech Republic

**Keywords:** Zoraptera, Polyneoptera, phylogeny, karyotype evolution, sex chromosomes

## Abstract

Our study provides the first comprehensive karyotype evolution analysis of the insect order Zoraptera. We present karyotypic descriptions of seven species across two families: Zorotypidae (*Usazoros hubbardi* and two *Zorotypus* spp.) and Spiralizoridae (*Centrozoros gurneyi*, *Spiralizoros magnicaudelli*, and two *Spiralizorose* spp.). These results facilitate a critical evaluation of existing cytogenetic knowledge in Zoraptera and the evolution of karyotypic traits across Polyneoptera. Most notably, we refute the presence of holocentric chromosomes in Zoraptera. Also, we show that the XY sex chromosome system is prevalent and likely ancestral within the order. Furthermore, by integrating the chromosome numbers of the studied species with a dated molecular phylogeny of Zoraptera, we provide the first estimation of the mode of chromosome number evolution for this group. Finally, standard karyotypic features (2n, chromosome morphology, and size) and the distribution of 18S rDNA and (TTAGG) telomeric motif clusters—detected by fluorescence in situ hybridization—reveal highly differentiated karyotypes and genomic structures. This genetic diversity contrasts sharply with the recognized morphological uniformity of Zoraptera.

## Introduction

Structural karyotypic changes have significant impact on organismal evolution by establishing inter-specific barriers (Faria & Navarro, 2010; Ortiz-Barrientos et al., 2016) and facilitating adaptation by spatial reorganization of genetic linkage groups (Guerrero & Kirkpatrick, 2014; Liu et al., 2022). However, modes of karyotype evolution may largely differ among evolutionary lineages, rendering immense variability in karyotype behavior and structure.

Insects, the most speciose animal class, exhibit remarkable variability in karyotypic features. This diversity includes fundamental differences in centromere organization (monocentricity/holocentricity), sex chromosome systems (SCSs), process of meiotic division and vast span of chromosome numbers, starting from 2n = 2 in *Myrmecia* ants (Crosland & Crozier, 1986; Debec et al., 2024) to 2n = 448–452 in *Polyommatus atlanticus* butterfly (Lukhtanov, 2015).

Variations in chromosome numbers are primarily driven by chromosomal fission, fusion and genome duplication (polyploidization). Recent advancements mathematical modeling of chromosome number evolution now allow for the estimation of structural change rates, providing a deeper characterization of karyotype evolution within specific lineages. By implementing these methods, specific modes of karyotype evolution have been detected in several insect orders (Alfieri et al., 2023; Sylvester et al., 2020). Polyneoptera, which comprises ten insect orders, represents one of the three primary evolutionary lineages of Neopterous insects, alongside Paraneoptera and Endopterygota (Misof et al., 2014). While characteristic modes of karyotype evolution have been identified in some polyneopteran groups—such as high rates of both fissions and fusions in Blattodea and a relatively high incidence of polyploidization in Phasmatodea (Sylvester et al., 2020)—data for other polyneopteran orders remain scarce or entirely absent.

Order Zoraptera, commonly known as angel insects, represents one of these data-deficient groups. For long time, their phylogenetic position remained elusive—a challenge famously termed the “Zoraptera problem” (Choe, 2018). Currently, Zoraptera are recognized as a sister lineage to the rest of Polyneoptera, either with Dermaptera (Wipfler et al., 2019) or alone (Tihelka et al., *preprint*). The group has been estimated to be of Late Paleozoic origin (Carboniferous-Permian) (Matsumura et al., 2020; Kočárková et al., 2026) and contains 47 currently recognized extant species (Kaláb et al., 2025) organized in two families (Spiralizoridae and Zorotypidae) and four subfamilies (Spiralizoridae: Latinozorinae, Spiralizorinae; Zorotypidae: Spermozorinae, Zorotypinae) (Kočárek et al., 2020). However, chromosome numbers are known only in four species (Jankásek et al., 2024; Kuznetsova et al., 2002).

First karyotype description of Zoraptera was performed in *Usazoros hubbardi* (2n♂ = 38) (Zoroypidae), suggesting neo-XY SCS and holocentric chromosomes to be present in the species (Kuznetsova et al., 2002). Consequently, in our previous study, we have described karyotypes of three Spiralizoridae species, expanding the span of chromosome numbers in Zoraptera to 2n♀ = 36–44 (Jankásek et al., 2024). Importantly, karyotypes of all of the studied species have been found to be monocentric. Also, both, X0 and XY SCSs have been observed in the studied species with the X chromosome being either telocentric (*Brazilozoros kukalovae*, X0 SCS) or metacentric (*B. huxleyi*, *Latinozoros cacaoensis,* XY SCSs).

In this study, we are newly presenting cytogenetic analyses of seven Zoraptera species (three from Zorotypidae, four from Spiralizoridae) including reevaluation of *U. hubbardi* karyotype and rejecting holocentricity in Zoraptera. Using the chromosome number counts and time calibrated phylogenetic tree, we were able to evaluate chromosome number evolution modes in ChromEvol v.3 software (Shafir et al., 2025). Moreover, we performed fluorescent *in situ* hybridizations (FISHs) to detect 18S rDNA and telomeric sequence clusters in all of the seven species. These sites represent standard cytogenetic markers. Clusters of rDNA are standard cytogenetic markers. Number and positions of 18S rDNA clusters generally illustrate dynamic or conservative evolution of genome structure (Šťáhlavský et al., 2018, 2020; respectively), while interstitial telomere sequences (ITSs) may highlight chromosome fusions (Vicari et al., 2022).

## Material and methods

### Sampling

For the time-calibrated molecular phylogeny analysis, 20 Zorapteran and 23 outgroups were utilized. The origin localities, DNA isolate codes and GenBank accession numbers for all DNA sequences are provided in Table S1. For karyotype analysis, we used samples from laboratory cultures obtained from individuals collected at the locations listed below. In Spiralizoridae we analyzed species: *Centrozoros gurneyi* (one female; Panama, Lagunas de Volcán, 8.7635N, 82.6782W); *Spiralizoros magnicaudelli* (two females; Malaysia: Brinchang Mt., 4.4594N, 101.3896E); *Spiralizoros* sp. (three females; Malaysia: Brinchang Mt., 4.4594N, 101.3896E); *Spiralizoros* sp. 3 (one male, Brunei: Ulu Temburong, 4.5596N, 115.1498E). In Zorotypidae we analyzed: *U. hubbardi* (three males and one female; USA: South Carolina, 34.71539N, 82.8411W); *Zorotypus* sp. 1 (one male, Madacascar, Analamazaotra, 18.9493S, 48.4269E); *Zorotypus* sp. 2 (three males, Madacascar, Ambohitantely, 18.1956S, 47.2878E). The living individuals were collected and reared following Jankásek et al. (2024).

### Chromosome slide preparations, imaging and karyotype descriptions

Seven Zoraptera species have been cytogenetically analyzed in this study. The abdomen cavity of analyzed individuals was hypotonized (0.075 M KCl, 35 min) and fixed in methanol:acetic acid solution (3:1, 20 min). Consequently, the abdomen content was dissolved in 60% acetic acid and spread in microscopic slides following the “plate spreading” method described by Traut (1976) and the chromosomes were stained for 10 min by 5% Giemsa-Romanowski solution in Sörensen phosphate buffer. The chromosome slides stained by Giemsa-Romanowski solution as well as slides subjected to FISH experiments were observed and photographed using an Olympus AX70 Provis microscope and Olympus DP72 camera with appropriate fluorescent filters.

The karyotypes were described using morphological chromosomal nomenclature established by Levan et al. (1964): metacentric (m), submetacentric (sm), subtelocentric (st), telocentric (t). The st and t chromosomes were pooled together into a st/t category. The chromosomes were measured in ImageJ 1.53p (Schneider et al., 2012) with Levan plugin (Sakamoto & Zacaro, 2009). The relative chromosomes length (RCL) was measured and calculated for the diploid set.

### 18S rDNA probe preparation

The 18S rDNA probe (∼1,800 bp, GenBank: OP558030.1) was amplified and DIG-11-dUTP labelled by PCR from genomic DNA of *B. kukalovae* using eukA: 5′-AACCTGGTTGATCCTGCCAGT3′ and eukB: 5′-TGATCCTTCTGCAGGTTCACCTACG-3′ primers (Medlin et al., 1988) and following PCR conditions: 95°C for 3 min, 35 cycles of 95 °C for 30 s, 51 °C for 40 s and 72 °C for 2 min. The final extension was at 72 °C for 10 min. Consequently, the PCR product was purified by Gel/PCR DNA Fragments Kit (Geneaid Biotech Ltd., New Taipei City, Taiwan) following the manufacturer’s protocol. The probe was mixed with competitive Salmon Sperm DNA and ethanol precipitated. The final probe mix contained 100ng of the probe and 25 µg of competitive DNA per slide in 50% formamide/2xSSC and 10% dextran sulphate.

### 18S rDNA and telomeric FISH

The 18S rDNA FISH was performed following the modified protocol of Fuková et al. (2005) as presented in Jankásek et al. (2024). Consequent stringency washes were done according to Sahara et al. (1999). Afterwards, each slide was incubated with 500 µl of 2.5% BSA /4xSSC blocking solution (20 min, room temperature (RT)). The probe was detected in 100 µl of anti-digoxigenin-fluorescein/2.5% BSA (10:100) solution for 1 h in RT and the slides were washed 3 times in 4xSSC/0.1% Tween (3 min, 37°C). At the end, the slides were counterstained with Mounting Medium with DAPI (Abcam plc., Cambridge, UK).

We used commercially prepared biotin-labelled (TTAGG)_8_ (Integrated DNA Technologies, Inc., Coralville, USA) probes to visualize telomere regions following the modified protocol of Cuadrado and Jouve (2010) as presented in Jankásek et al. (2024).

To account for relative positions of telomeric and 18S rDNA loci, individual slides were reprobed and photographed after each experiment. After imaging, the results of the first FISH experiment (either telomeric or 18S rDNA), the cover glass was gently removed and the slides were washed in 4xSSC/0.1%Tween (shaking, 30 min, RT). Subsequently the slides went through an ethanol series (70%, 80%, 96%, 2 min each), were fixed in methanol:acetic acid (3:1) solution (30 min, RT), air dried and aged in 60°C on histological plate for 1 hour and the slides were stored.

Right before reprobing in the subsequent FISH experiment, the slides were washed in 2xSSC (5 min, RT), fixed in 4% paraformaldehyde/1xPBS (5 min, RT) and washed again in 2xSSC (5 min, RT). After, the old probe was removed by strong stringency washing in 70% formamide/1% TritonX/0.1xSSC (15 min, 70°C), slides went through an ethanol row (-20 °C 70%, 80%, 96%, 2 min each) and new probe was applied according to the protocols above.

### DNA sample collection and editing

The time calibrated Bayesian inference (BI) phylogenetic analysis was based on sequences of 12S rRNA, 16S rRNA, 18S rRNA, 28S rRNA, H3 and COI genes from 43 species (20 Zoraptera, 23 outgroups). Most of the sequences were downloaded from GenBank (Sayers et al., 2025) except for *Spiralizoros* sp., *Zorotypus* sp. 1 and *Zorotypus* sp. 2 whose sequences have been generated in this study. DNA of these species was isolated from leg muscle tissue using Tissue DNA kit, genomic DNA isolation from tissue, E.Z.N.A.® (Avantor, Inc., Radnor, USA) and following the manufacturer’s protocol. List of used primers, PCR protocols and respective protocols are given in Table S2. The amplicons were purified using Gel/PCR DNA Fragments Extraction Kit (Geneaid Biotech Ltd., New Taipei, Taiwan), following the manufacturer’s protocol, and Sanger sequenced (Laboratory OMICS – Genomics, Biocev, Czech Republic). Sequence assembly, editing and concatenation was performed in Geneious 11.15 (Kearse et al., 2012). Sequences of each gene were aligned in MAFFT v7 web application (Katoh et al., 2017) using Q-INS-i iterative refinement method. The alignments of rRNA gene sequences were quality filtered using Gblocks 0.91b (Lemoine et al., 2019). The final concatenated gene matrix was 4 537 nt long.

### Phylogenetic Bayesian inference analysis

The BI phylogenetic analysis was performed in BEAST2 2.7.7 (Bouckaert et al., 2014). Individual substitution models estimated by BEAST Model Test (Bouckaert & Drummond, 2017) package were assigned to each of the rRNA gene fragments and to each of the codon positions in coding genes (COI, H3). Uncorrelated relaxed lognormal model was used for estimation of common molecular clock rate for all the genes. The molecular clock calibration points were set following Kočárková et al. (2026). The tree prior was set to birth death model. Two 150 million generations long MCMC chains with each 5^th^ thousand generation being saved were run. The estimated trees were combined with 25% burn-in using LogCombiner 2.7.7 and the resulting tree was annotated in TreeAnnotator 2.7.7.

### Reconstruction of chromosome number evolution

The estimation of chromosome number evolution mode and ancestral chromosome numbers by BI, marginal reconstruction (MR) maximum likelihood (ML) was performed using ChromEvol v.3 (Shafir et al., 2025). Since polyploidization is considered rather rare in insects (Lokki & Saura, 1980; Nakatani & McLysaght, 2019), it was not considered in the estimation of chromosome number evolution model in Zoraptera. The minimum and maximum allowed haploid chromosome numbers were set to 6 and 32. The optimal functions to model chromosome fission and fusion rates were evaluated following Shafir et al. (2025). Each of the optimization runs had 10 thousand simulations. The optimal modelling functions were chosen based on their AIC scores. The optimized run had 100 million simulations and allowed for transitions between chromosome number evolution models within the time calibrated phylogenetic tree.

## Results

Generally, obtaining sufficient number of good quality metaphase images was found to be excessively challenging in Zoraptera. Out of the seven studied species, accurate chromosome number could not be detected in *Zorotypus* sp. 2. Quality of obtained figures did not allow us to estimate chromosome morphology in *Zorotypus* sp. 1 and *Zorotypus* sp. 2. Also, reprobing with 18S rDNA probe did not produce any conclusive signal in *Spiralizoros* sp. 3. Results of cytogenetic and ancestral chromosome number estimation analyses are summarized in Fig. 9.

### Spiralozoridae

#### Centrozoros gurneyi (Choe, 1989)

This parthenogenetic population exhibited 2n♀ = 36 (Fig. 1). The largest chromosome pair was sm and represented approx. 7.3 % of RCL. Second and third largest chromosome pairs were m and represented 5.0 and 4.0 % of RCL, respectively. The rest of the chromosomes were st/t and their relative size was gradually decreasing from approx. 3.27 to 1.18 % of RCL.

**Fig. 1:**
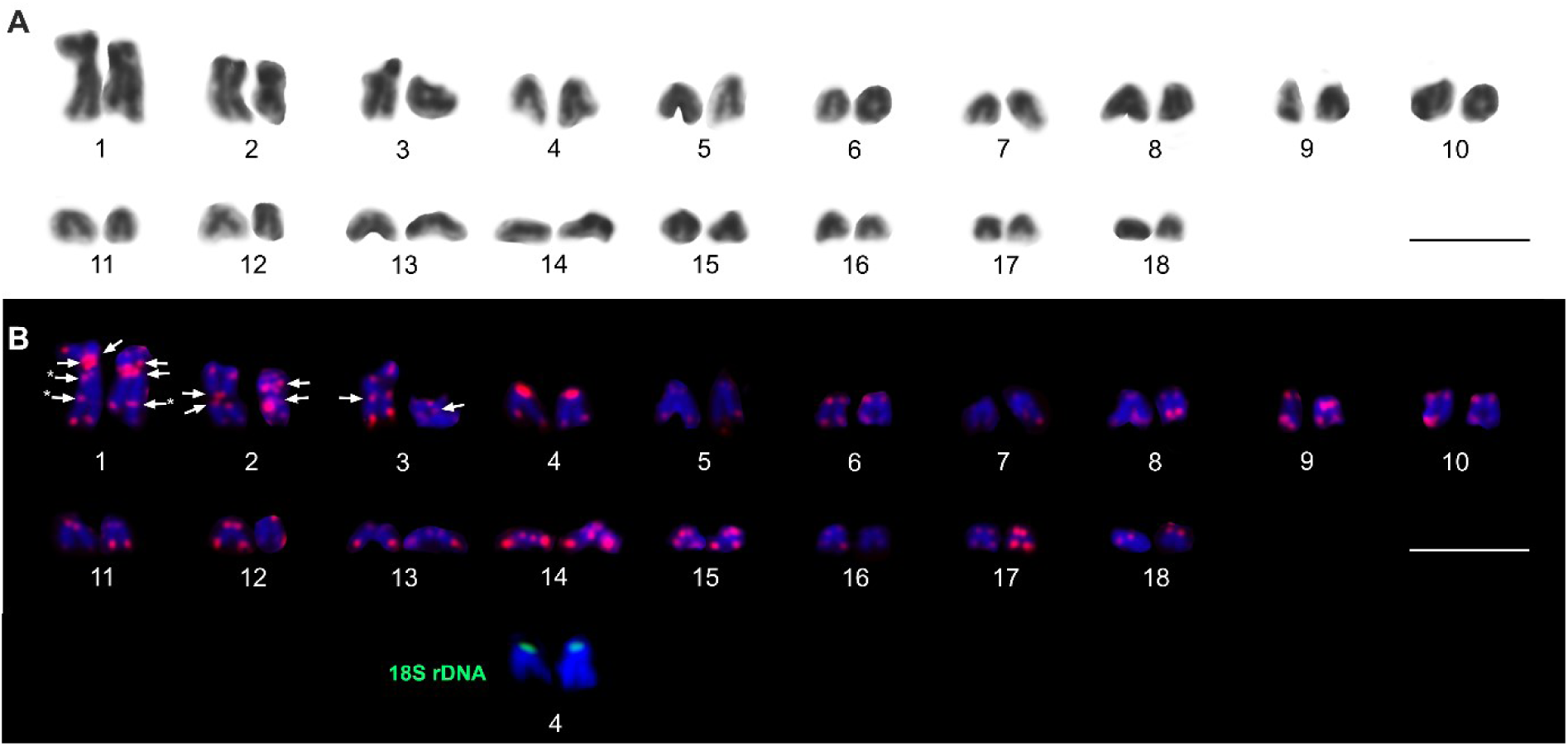
Female karyotype of Centrozoros gurneyi. A: Chromosomes after Giemsa staining. B: Chromosomes after (TTAGG)8 (red signals) and 18S rDNA (green signals) FISH counterstained with DAPI. The arrows highlight interstitial telomeric sequences. The arrows with asterisk highlight telomeric signals from overlapping chromosomes. Scale bars = 10 μm.

The FISH with 18S rDNA probe revealed one pair of terminal clusters on p arms of fourth chromosome pair (st/t). The FISH with (TTAGG)_8_ probe revealed standard telomeric pattern. Additionally, ITSs were revealed in all of the three largest chromosome pairs. Two pericentric ITSs, on p and q arms, have been detected in the largest chromosome pair. Two pericentric ITSs, were located on p and q arms of the second largest m chromosome and one pericentric ITS was located on the third largest m chromosome (Fig. 1B).

#### *Spiralizoros magnicaudelli* (Mashimo, Engel, Dallai, Beutel & Machida, 2013)

The species exhibited 2n♀ = 44 chromosomes with most of them being clearly metacentric or submetacentric (Fig. 2). The largest two chromosome pairs are sm and represents approx. 4.8 % and 3.92 % of RCL.

**Fig. 2:**
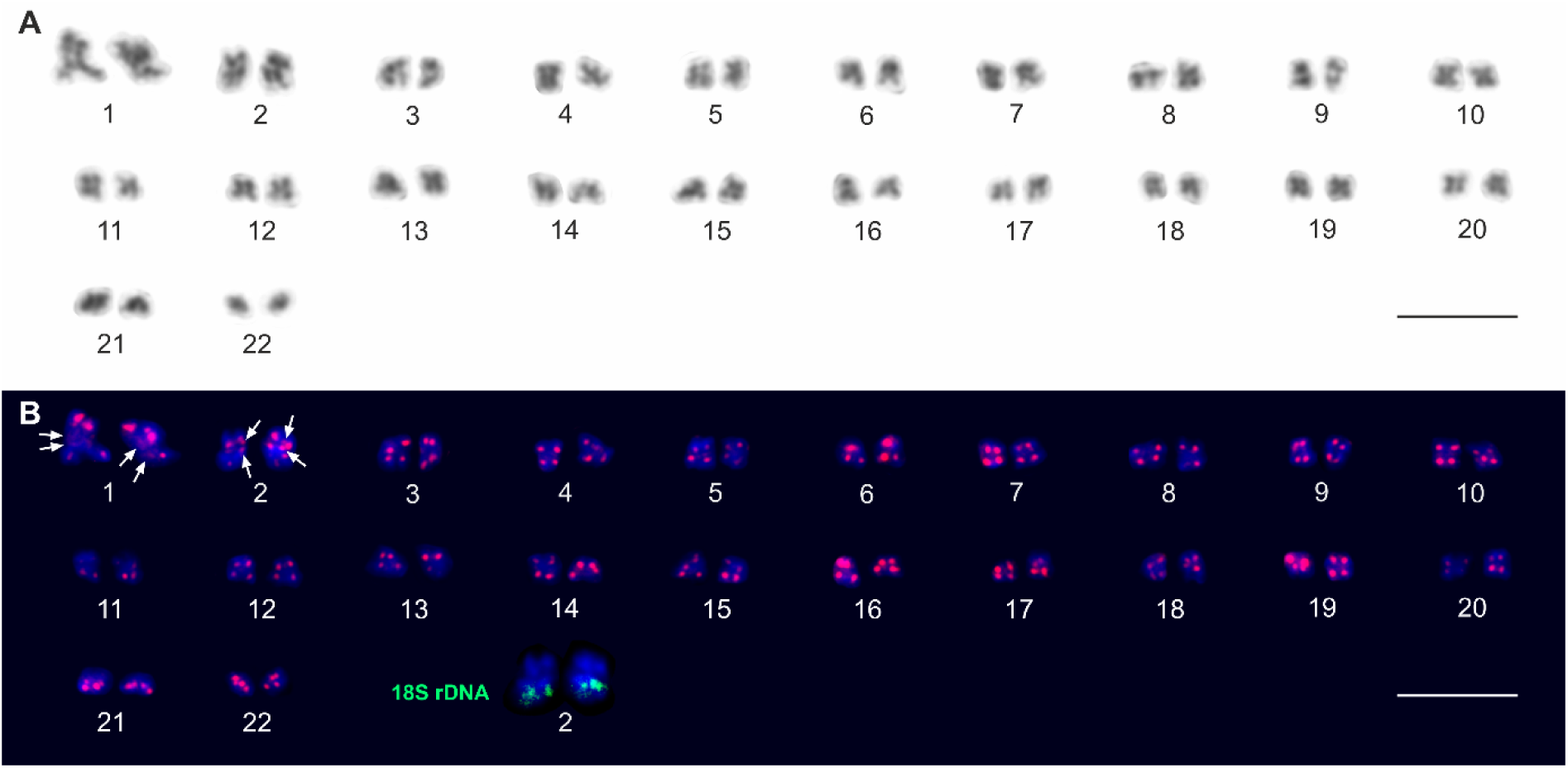
Female karyotype of Spiralizoros magnicaudelli. A: Chromosomes after Giemsa staining. B: Chromosomes after (TTAGG)8 (red signals) and 18S rDNA (green signals) FISH counterstained with DAPI. The arrows highlight interstitial telomeric sequences. Scale bars = 10 μm.

The FISH with 18S rDNA probe visualized one pair of subterminal clusters, each on a q arm of second largest chromosome pair (Fig. 2B). The FISH with (TTAGG)_8_ probe showed standard pattern with almost all telomeric regions being visualized. Additionally, dispersed pericentric signals were observed on both centromere sides on each chromosome of the largest pair. Also, two pericentric ITSs, each on one chromosome arm, were visualized on each of the second largest chromosome pair (Fig. 2B).

#### Spiralizoros sp

The species exhibited 2n♀ = 40 (Fig. 3). The two largest chromosomes represent heteromorphic pair composed of sm and m chromosomes representing approx. 6.0 % and 5.4 % of RCL. Noteworthy, the q arm of the smaller chromosome from this pair was positively heteropycnotic (Fig. 3B). The length difference between the two chromosomes was apparently smaller during pachytene (Fig. 7).

**Fig. 3:**
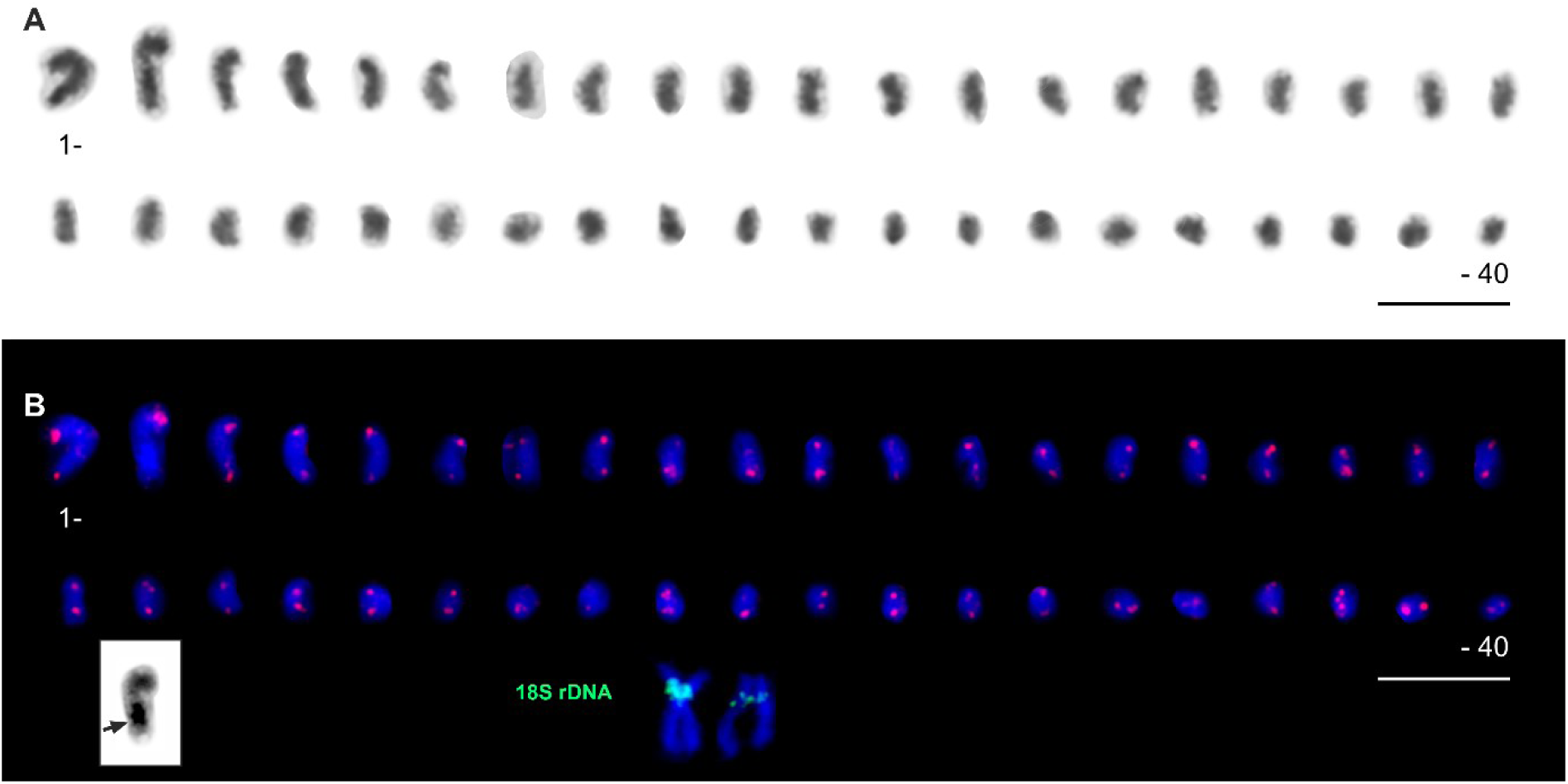
Female karyotype of Spiralizoros sp. A: Mitotic metaphase chromosomes after Giemsa staining. B: Mitotic metaphase chromosomes after (TTAGG)8 (red signals) and 18S rDNA (green signals) FISH counterstained with DAPI. The black arrow in the inset highlights positively heteropycnotic region in the second largest chromosome. Scale bars = 10 μm.

The FISH with 18S rDNA probe visualized one pair of pericentric clusters on p arms of a heteromorphic chromosome pair. One of the clusters is larger and localized on a metacentric whereas the other cluster is on sm chromosome (Fig. 3B). Beside revealing standard telomeric pattern, the FISH with (TTAGG)_8_ probe showed two ITS clusters on q arms of the two largest chromosomes (Fig. 7A).

#### Spiralizoros sp. 3

The species exhibited 2n♂ = 42 (Fig. 4). The poor spiralization of chromosomes did not allow for detailed specification of chromosome morphology. However, it is recognizable that most of the chromosomes are st/t. The two largest chromosomes are clearly st/t and represented 6.9 and 5.6 % of RCL. The length of remaining chromosomes gradually decreased from 5.2 to 1.0 % of RCL. From third to tenth largest chromosome the p arms were positively heteropycnotic (Fig. 3A)

**Fig. 4:**
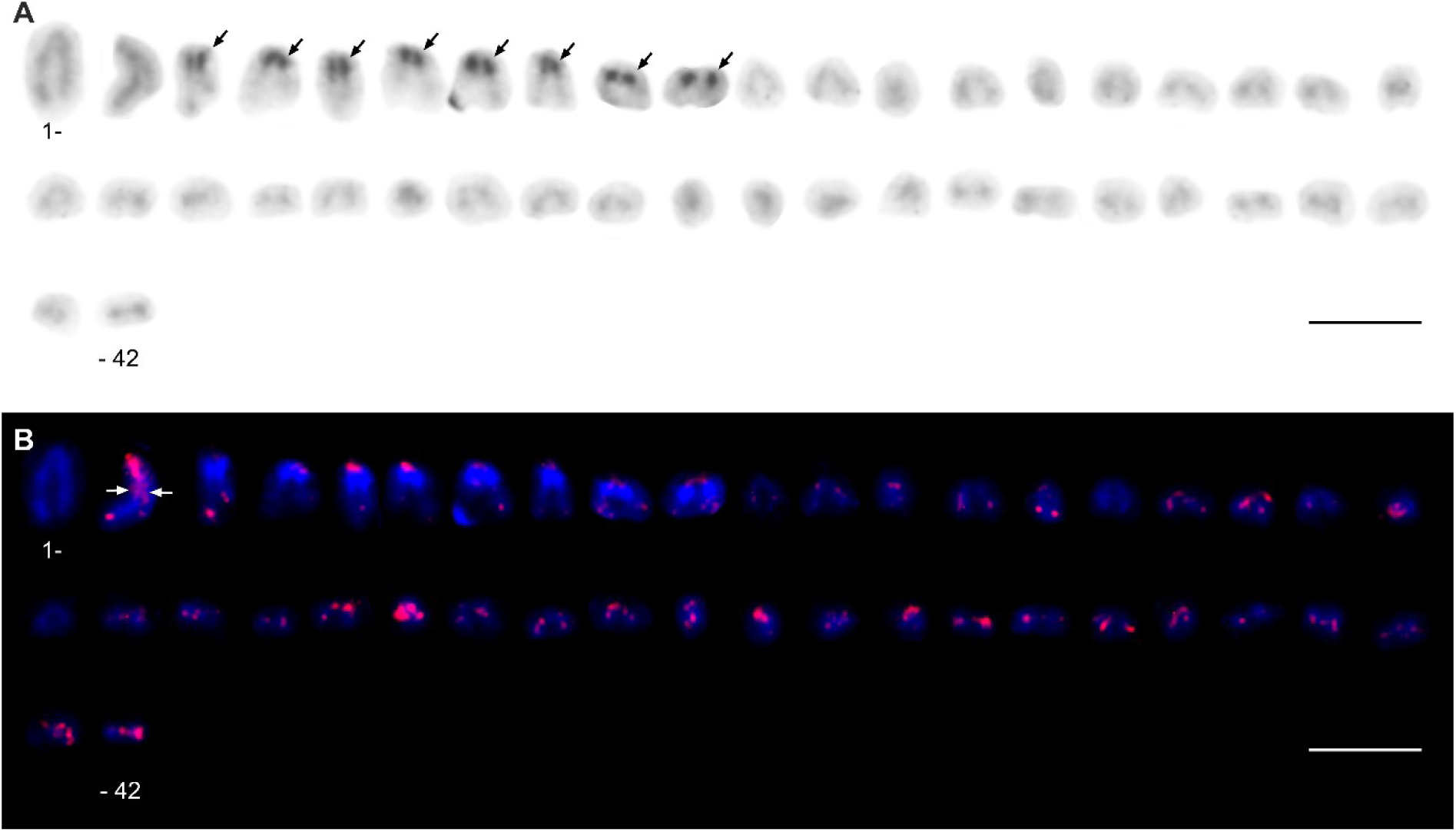
Male karyotype of Spiralizoros sp. 3. A: chromosomes after Giemsa staining. B: Chromosomes after (TTAGG)8 FISH (red signals) counterstained with DAPI. The black arrows highlight positively heteropycnotic p arms of chromosome 3 to 10. The white arrow highlights interspersed interstitial telomeric sequences. Scale bars = 10 μm.

The FISH with 18S rDNA probe did not produce any conclusive results. The FISH with (TTAGG)_8_ showed standard telomeric pattern with additional dispersed ITS signals throughout the length of the second largest chromosome (Fig. 4B).

#### Usazoros hubbardi (Caudell, 1918)

The species exhibited 2n♂, ♀ = 38, 38 (Fig. 5). In males, the two largest chromosomes were m and sm and represented 9.0 and 7.3 % of RCL and the rest of chromosomes gradually decreased from 6 to 3.9 %. The quality of metaphase figures did not allow for detailed chromosomal morphology measurements, however, most of the chromosomes are either m or sm.

**Fig. 5:**
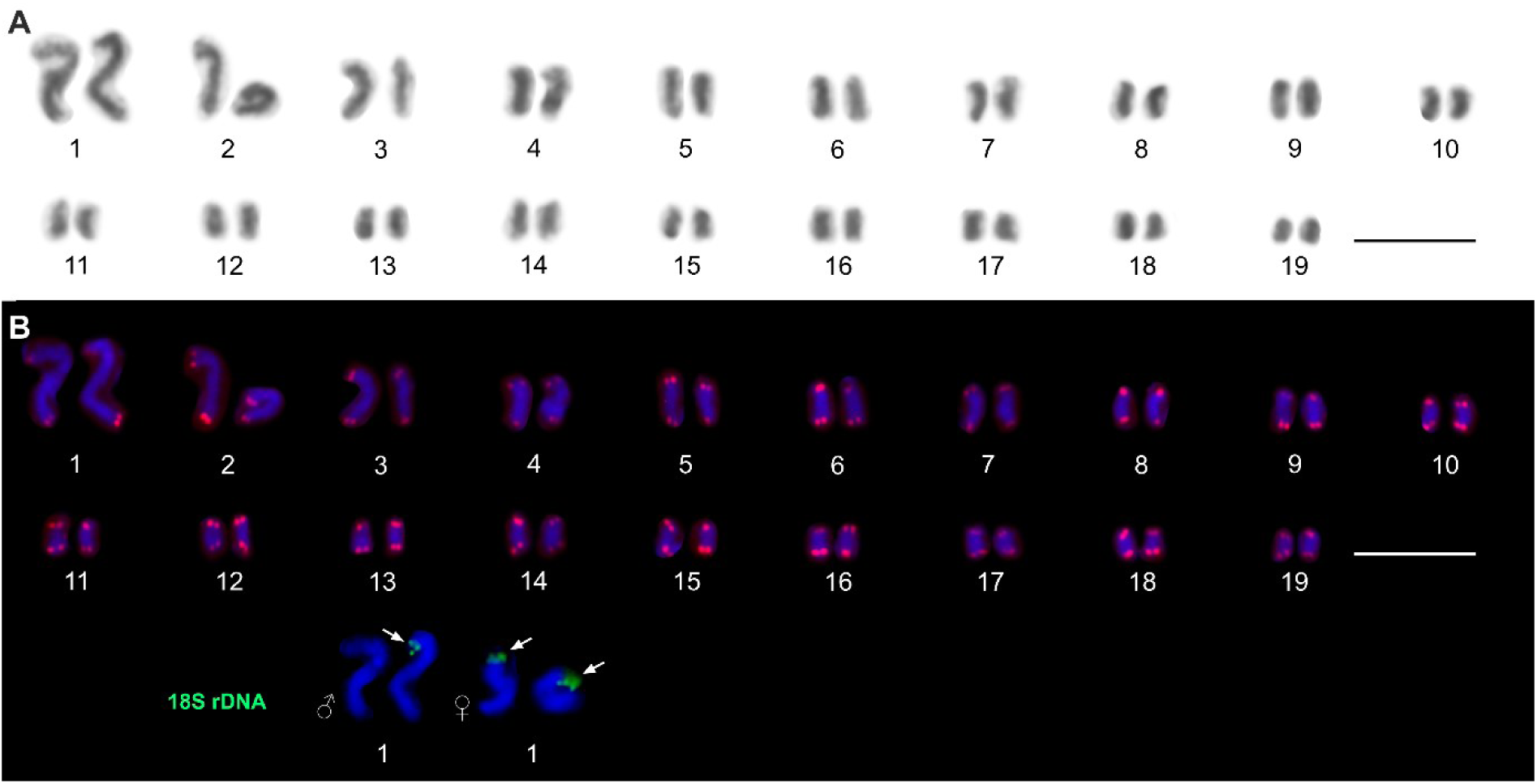
Male karyotype of Usazoros hubbardi. A: Mitotic metaphase chromosomes after Giemsa staining. B: Mitotic metaphase after (TTAGG)8 FISH (red signals) counterstained with DAPI. Both, male and female first chromosome pair after 18S rDNA FISH (green signals highlighted by white arrows) counterstained with DAPI is also presented. Scale bars = 10 μm.

The FISH with 18S rDNA probe revealed one terminal cluster on the q arm of the largest chromosome in males (three males studied) (Fig. 5B). In female (one female studied), one pair of 18S rDNA clusters was observed, each of the clusters being localized terminally on q arms of the largest chromosome pair (Fig. 5B). The FISH with (TTAGG)_8_ revealed standard telomeric pattern without ITSs (Fig. 5B).

#### Zorotypus sp. 1

Only two sister male metaphase II figures were available for karyotype description of the species. Each of the metaphases II had n = 11 (Fig. 7B-G), assuming the 2n = 22 (Fig. 6). The quality of the figures did not allow for detailed chromosomal morphology measurements. The two largest chromosomes represented 7.29 % and 6.47 % of RCL which was gradually decreasing from 5.22 % to 3.74 % in remaining chromosomes. The last two smallest chromosomes represented 2.58 % and 2.43 % of RCL.

**Fig. 6:**
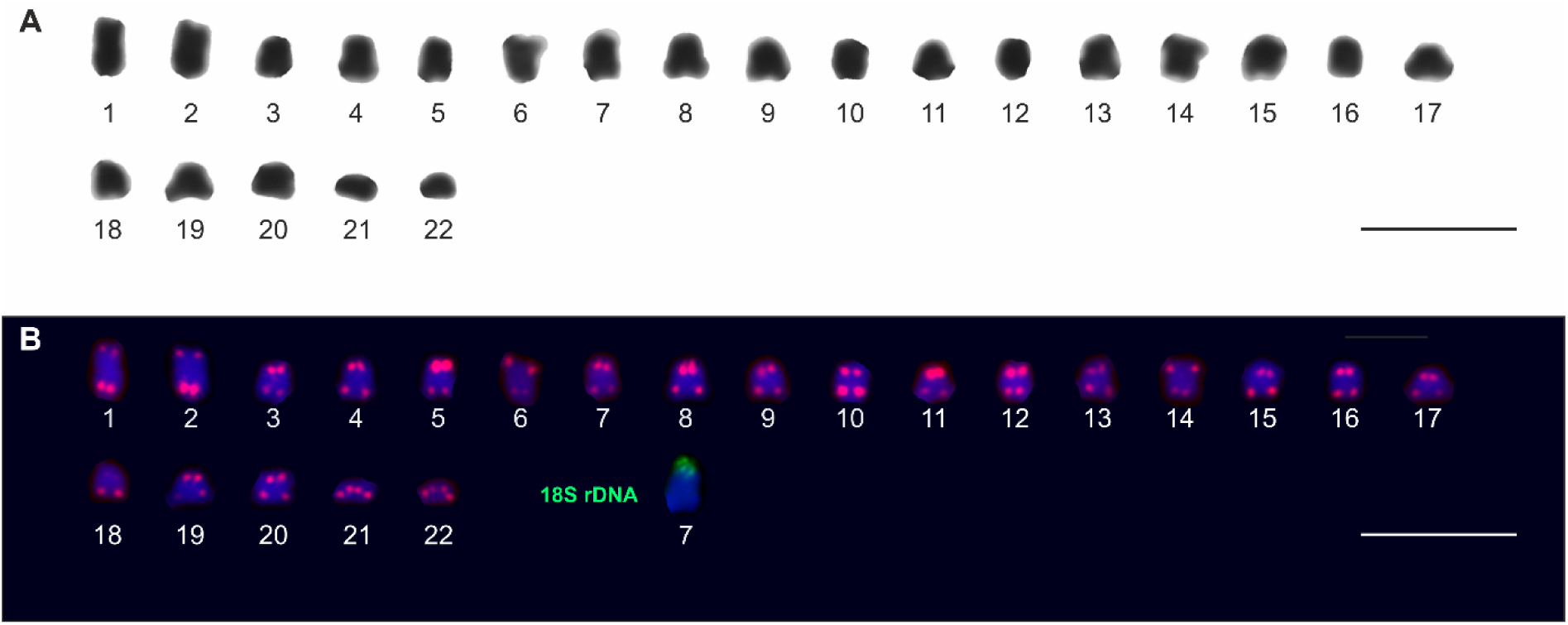
Male karyotype of Zorotypus sp. 1. A: Chromosomes compiled from two sister metaphases II after Giemsa staining. B: Chromosomes compiled from two sister metaphases II after (TTAGG)8 (red signals) and 18S rDNA (green signal) FISH counterstained with DAPI. Scale bars = 10 μm.

**Fig. 7:**
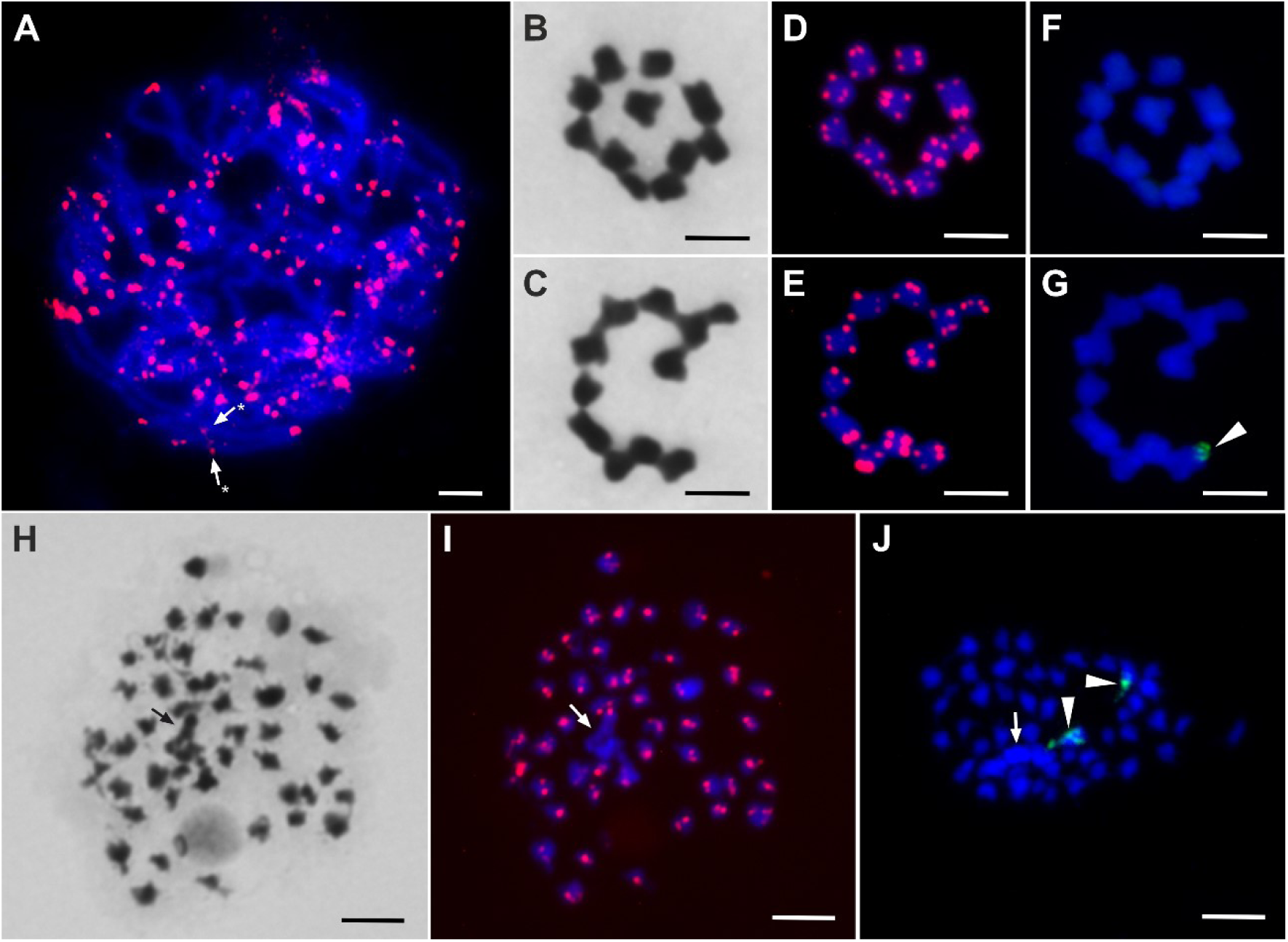
A: Spiralizoros sp. female pachytene of after (TTAGG)8 FISH (red signals) counterstained with DAPI, arrows with asterisks highlight interstitial telomeric sequences in the largest chromosome pair in the female karyotype. B, C: Male Zorotypus sp. 1 sister metaphases II (n = 11) after Giemsa staining. D, E: The same metaphases II after (TTAGG)8 FISH (red signals) counterstained with DAPI. F, G: The same metaphases II after 18S rDNA FISH (green signal highlighted by arrowhead) counterstained with DAPI. H: Zorotypus sp. 2 male mitotic prophase after Giemsa staining, the black arrow highlights ambiguous chromatin accumulation. I: The same prophase after (TTAGG)8 FISH (red signals) counterstained with DAPI, the white arrow highlights the same ambiguous chromatin accumulation. J: Another Zorotypus sp. 2 male mitotic prophase after 18S rDNA FISH (green signals highlighted by arrowheads), the arrow highlights the same ambiguous chromatin accumulation. Scale bars = 5 μm.

The FISH with 18S rDNA probe revealed one terminal cluster in only one of the two sister haploid sets, located on one of the medium sized chromosomes (Fig. 6B; 7G). The FISH with (TTAGG)_8_ probe revealed standard telomeric pattern in both of the studied metaphases II (Fig. 6B; 7D, E).

#### Zorotypus sp. 2

The species exhibited approximately 2n♂ = ∼44 (Fig. 7H-J). Only male mitotic prophases with advanced spiralization have been obtained in the species. The exact chromosome number could not be determined since ambiguous chromatin accumulation is formed around a large chromosome in the obtained male prophases.

The FISH with 18S rDNA probe revealed two clusters, each on different chromosome (Fig. 7J). The FISH with (TTAGG)_8_ probe reveled signals on each of the chromosomes (Fig. 7I). However, poor chromatin spiralization did not allow for precise localization of the 18S rDNA and telomeric signals.

The full time calibrated phylogenetic tree with all outgroup species is presented in Fig. S1. The result tree focused on Zoraptera is presented in Fig. 8. The origin of recent Zoraptera and split between Spiralizoridae and Zorotypidae is estimated to 302.14 Mya (95% CI = 241.61–361.22 Mya). The tree is organized into two main evolutionary lineages – families Zorotypidae and Spiralizoridae. The origin of Zorotypidae was estimated to 241.03 Mya (95% CI = 173.83 – 305.64 Mya, pp = 1). The family was further split into two clades, representing subfamilies Spermozorinae (genus *Spermozoros*, pp = 1) and Zorotypinae (genera *Usazoros* and *Zorotypus*, pp = 1). The genus *Spermozoros* was estimated to emerge 117.71 Mya (95% CI = 52.70–189.45 Mya). The split between *Usazoros hubbardi* lineage and *Zorotypus* was estimated to 182.46 Mya (95% CI = 114.30–250.62 Mya). Spiralizoridae origin was estimated to 226.48 Mya (95% CI = 161.79–291.36 Mya, calibration point). The family was also divided into two clades representing subfamilies Latinozorinae (genus *Latinozoros*) and Spiralizorinae (genera *Brazilozoros*, *Centrozoros*, *Scapulizoros*, *Spiralizoros*). The split between the subfamilies was estimated to 226.48 Mya (95% CI = 161.79–291.36 Mya). Spiralizorinae was further divided into sister clades (*Scapulizoros* + *Spiralizoros*) with estimated origin 144.78 Mya (95% CI = 93.62–198.26, pp = 0.62) and (*Brazilozoros* + *Centrozoros*) with estimated origin 120.46 Mya (95% CI = 66.34–177.05 Mya, calibration point).

**Fig. 8:**
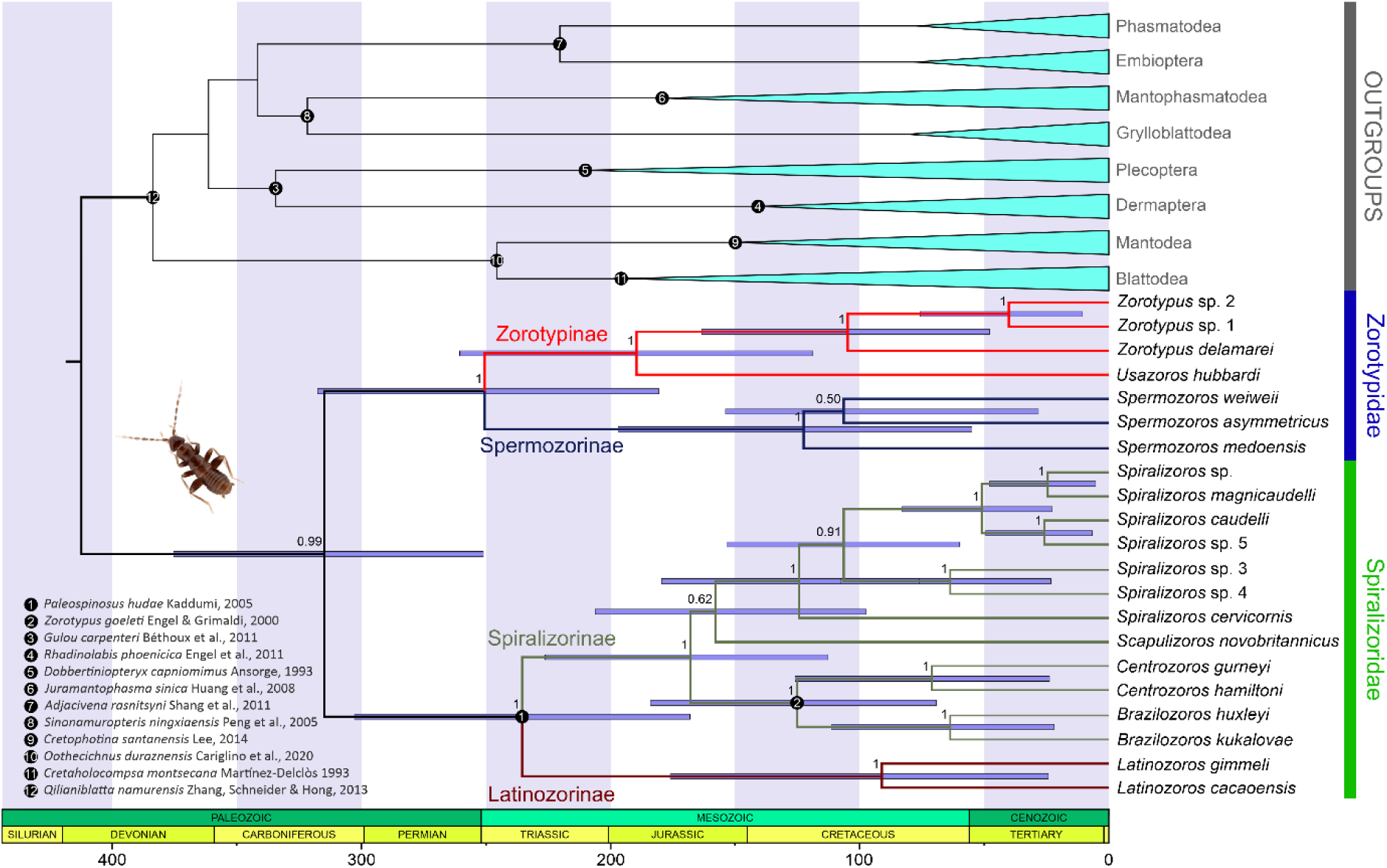
Time calibrated phylogenetic tree of Zoraptera produced by Bayesian inference method. Black circles with numbers represent fossil calibration points. Posterior probability is represented by the numbers next to the tree nodes. The blue bars around the tree nodes represent 95% confidence intervals.

**Fig. 9:**
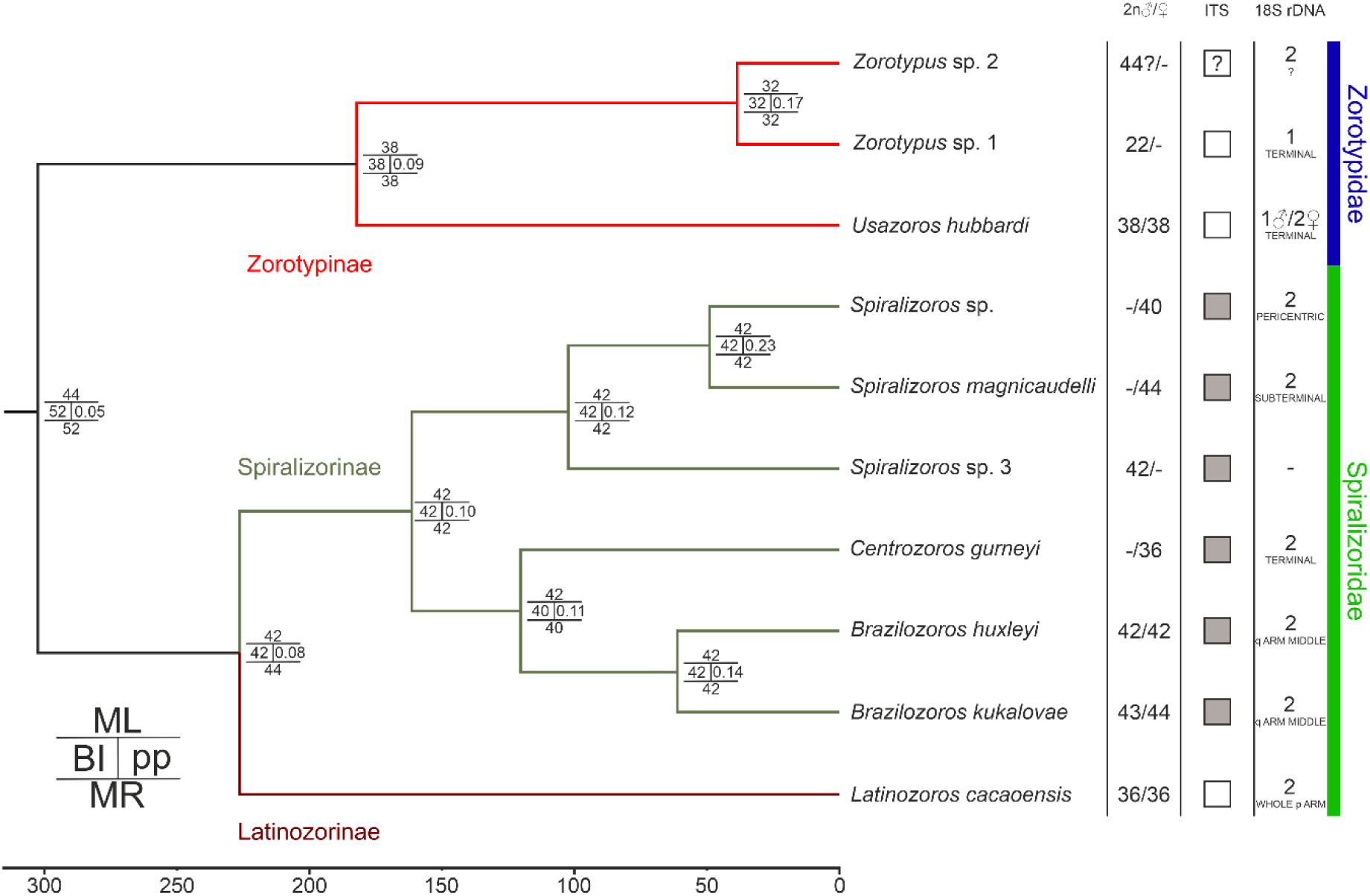
Time calibrated Bayesian inference constructed phylogenetic tree of Zoraptera with species mapped male and/or female diploid chromosome numbers, presence or absence of interstitial telomeric sequence (ITS) and number of 18S rDNA and their localization. The matrices next to the phylogenetic tree nodes represent ChromEvol3 estimated ancestral diploid chromosome numbers by maximum likelihood (ML), Bayesian inference (BI) and marginal reconstruction (MR) methods.

### Chromosome number evolution

A single chromosome number evolution model was estimated for the whole Zoraptera. Based on AIC scores, linear and univariate linear functions were estimated as the best fit models for chromosome fission and fusion rates, respectively. The estimated function of chromosome fusion rate was:

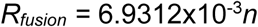

where *R_fusion_* is the chromosome fusion rate per species per million years and *n* is the default haploid chromosome number before the fusion. The chromosome fission rate was given by the function:

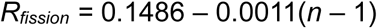

Where *R_fission_* is the chromosome fission rate per species per million years and *n* is the default haploid chromosome number before the fission. This means that, in both types of rearrangements, rates of change from *n* to a chromosome number larger (fission) or smaller (fusion) by one are dependent on the default haploid chromosome number. The estimated ancestral chromosome numbers (in diploid set) by the chromosome evolution modes are presented in Fig. 9. The ancestral 2n = 44 (ML) or 2n = 52 (BI, MR) were estimated for the whole Zoraptera albeit with low support (pp = 0.05). Generally, supports for ancestral chromosome numbers in Zorotypinae (2n = 32; ML, BI, MR) and Spiralizoridae (2n = 42; ML, BI, MR) were also relatively low (pp = 0.09 and 0.08, respectively). While the 2n = 42 was found to be relatively stable in Spiralizoridae, significant changes in chromosome numbers in the Malagasy *Zorotypus* spp. have been estimated in Zorotypinae (Fig. 9).

## Discussion

### Cytogenetic characteristics of Zoraptera

The cytogenetically examined species manifest 2n = 22–44. Chromosome number has been previously detected only in *U. hubbardi* (2n♂ = 38) (Kuznetsova et al., 2002). Our results show that both sexes of the species share the same chromosome number (2n♂, ♀ = 38, 38). Consistent with Jankásek et al. (2024), all cytogenetically examined Zoraptera species in this study possess monocentric chromosomes. Consequently, we reject the suggestion of Kuznetsova et al. (2002), that Zoraptera possess holocentric chromosomes, as well as that holocentricity is a synapomorphy of Dermaptera and Zoraptera (Kuznetsova et al., 2002). Despite generally poor spiralization of obtained metaphase chromosomes, it is obvious that both chromosome number and morphology is changing in Zoraptera evolution, meaning the group is not in karyotype evolution stasis. This is especially apparent in *Spiralizoros* genus where karyotypes of *Spiralizoros* sp. 3 and *S. magnicaudelli* substantially differ in prevailing chromosome morphology types.

The FISH experiments with (TTAGG)_8_ revealed telomeric signals in all cytogenetically studied species. However, we have shown in our previous study that FISH with both (TTAGG)_8_ and (TTAGGG)_8_ probes render identical signals in Zoraptera using low stringency washing FISH protocol (Jankásek et al., 2024). Both of the probe types did not render conclusive signals when stronger stringency washing was applied. Also, single-primer PCR experiments with (TTAGG)_6_ and (TTAGGG)_6_ primers did not render positive results in Zoraptera, meaning none of the two motifs form telomeres in this insect order (provided in *Review History* section of Jankásek et al., 2024). This means that our FISH experiments do not identify the unknown Zorapteran telomeric motif (Kuznetsova et al., 2020), but show that it is sufficiently similar in sequence. The absence of (TTAGG) canonic telomeric motifs has been also shown in several other insect orders (Frydrychová et al., 2004). Interestingly, in Dermaptera, an order closely related to Zoraptera (Wipfler et al., 2019; but see Tihelka et al., *preprint*), TTCGG telomeric motif has been recently identified (Lukhtanov, 2025). The standard distribution of telomeric signals without ITSs in *Zorotypus* sp. 1, *U. hubbardi* (Zorotypinae, Zorotypidae) (present study) and *L. cacaoensis* (Latinozorinae, Spiralizoridae) (Jankásek et al., 2024) is in striking difference with the presence of ITSs in all of the so far cytogenetically studied representatives of Spiralizorinae (Spiralizoridae) (Jankásek et al., 2024; this study). ITSs are generally rare in insects (Vicari et al., 2022), however, they have been identified in Lepidoptera (Rego & Marec, 2003), Hemiptera (Chirino et al., 2017) and, most commonly, in Orthoptera (reviewed in Vicari et al., 2022). In some cases, the ITSs represent relics after chromosome fusions (reviewed in Vicari et al., 2022). The mostly pericentric ITSs in Spiralizorinae (Jankásek et al., 2024; this study) may be relics from Robertsonian fusions of st/t chromosomes, forming relatively large sm and m chromosomes. However, this would ideally need to be tested by comparative genomic studies since other mechanisms like inversions, gene conversion, illegitimate homologous (ectopic) recombination or telomerase activity during non-homologous end joining (NHEJ) double strand repair process may also generate ITSs (Aksenova & Mirkin, 2019; Slijepcevic, 1998; Vicari et al., 2022). Consequently, their enlargement may be facilitated by unequal recombination or DNA polymerase slippage (Aksenova & Mirkin, 2019).

However, basic FISH detectable ITSs are generally not common and mostly do not accompany chromosome fusion points, probably due to their effective elimination after the fusion. The deleterious effect of ITSs, as well as its evolutionary potential, resides in increasing the recombination rate and consequent structural instability in the given genomic region; the ITSs act as chromosome rearrangement hot spots (Aksenova & Mirkin, 2019; Rosas Bringas et al., 2024). In Spiralizorinae, the proximal location to centromeres may shelter the ITSs since (peri)centric regions are generally composed of heterochromatin and have low recombination rates (Nambiar & Smith, 2016), protecting the ITSs from recombination induced deletions.

Inter-specific variation in 18S rDNA clusters’ position also manifested dynamic structural evolution of Zoraptera genome. In all but one of the so far cytogenetically analyzed Zoraptera species, two 18S rDNA clusters have been detected, corresponding to the median value of 45S rDNA sites (consisting of 18S rDNA, 5.8S rDNA, 28S rDNA and two external and two internal transcribed spacers) in animals (Sochorová et al., 2018). The 18S rDNA clusters in (sub)terminal (*C. gurneyi*, *S. magnicaudelli*, *U. hubbardi*, *Zorotypus* sp. 1) and pericentric regions (*Spiralizoros* sp.) as well as pericentric to terminal on p arms (*L. cacaoensis* (Jankásek et al., 2024)) are the most common in arthropods. Contrary to that, interstitial 18S rDNA clusters detected in *B. huxleyi* and *B. kukalovae* are relatively rare in the group (Sochorová et al., 2018). It has been hypothesized that terminal localization of rDNA is common probably due to generally more frequent recombination in distal chromosome regions, facilitating the rDNA’s typical concerted evolution mode and ectopic recombination (reviewed in Sochorová et al., 2018). On the other hand, the common pericentric localization probably associates with rDNA’s facilitation of heterochromatin formation (often called “position effect”) and securing the centromere function (reviewed in Sochorová et al., 2018). Congruently, the subterminal 18S rDNA cluster in *S. magnicaudelli* is present in a very prominent secondary constriction (Fig. 5), which is probably formed due to the rDNA’s “position effect”. Although fragmentary, the available data allow us to propose that interstitial 18S rDNA clusters may be specific for *Brazilozoros* genus. This exact 18S rDNA cluster position has not been detected in any other species cytogenetically studied species.

Moreover, females and males of *U. hubbardi* have been shown to differ in 18S rDNA cluster number (Fig. 5). Only one terminal cluster on the one of the two largest chromosomes has been detected in males, but two such clusters have been detected in a female. Based on these results and meiosis observations of Kuznetsova et al. (2002), we identify that the chromosome bearing the cluster is an X chromosome and we confirm that *U. hubbardi* has XY SCS (Kuznetsova et al., 2002). In Zorotypinae (Zorotypidae), one terminal 18S rDNA cluster has also been detected in one of the sister male metaphases II of Malagasy *Zorotypus* sp. 1 (2n♂ = 22). Although a female karyotype of the species has not been studied here, we hypothesize the 18S rDNA cluster may also serve as an X chromosome marker in the species, possibly indicating the XY SCS. Interestingly, *Zorotypus* sp. 2, the other closely related Malagasy species, has two 18S rDNA clusters and highly contrasting chromosome number (2n♂ =∼44), suggesting substantially divergent karyotype evolution between the two closely related species. In the male karyotype of *Spiralizoros* sp. 3, the second largest chromosome with dispersed ITSs may also be a sex chromosome though, this needs to be confirmed by analysis of a female karyotype. In female karyotype of *Spiralizoros* sp., the two largest and heteromorphic chromosomes bear an ITS on q arms. The heteromorphy is caused by stronger spiralization of one of the chromosomes, which is weaker in pachytene, when the size difference between the chromosomes is not so prominent (Fig. 7A). The difference in spiralization level between homologous chromosomes in a female may be due to dosage compensation of one of the putative X chromosomes. Our results suggest XY SCS is dominant in Zoraptera and probably ancestral. In Dermaptera, an order considered sister to Zoraptera (Wipfler et al., 2019; but see Kočárková et al., 2026; Tihelka et al, *preprint*), the XY SCS is ancestral (White, 1976). Since both of the insect orders represent earliest branching Polyneopteran lineages (Wipfler et al., 2019; Tihelka et al, *preprint*), this raises a question whether an XY SCS is ancestral in Polyneoptera and the X0 SCS ancestral in most of the other Polyneopteran orders was secondarily derived by loss of the Y chromosome. However, to study this, the homology between the Dermapteran and Zorapteran XY SCSs would have to be confirmed.

### Phylogeny

The result phylogenetic topology of the BI analysis is largely congruent with previous results of Kočárková et al. (2026) and Matsumura et al. (2020). In addition, our analysis also provides phylogenetic positions of previously unanalyzed Malagasy *Zorotypus* spp. and one Malayan *Spiralizoros* sp. The estimated origins of recent Zoraptera and Spiralizoridae and Zorotypidae families is congruent with the results of Kočárková et al. (2026). Our estimated divergence times on species and genera level are generally older than presented by Kočárková et al. (*2026*). However, the 95% CIs estimated herein and in previous studies largely overlap (Matsumura et al., 2020; Kočárková et al., 2026). Recent interest in Zoraptera and broader sampling of more species and loci will allow for more precise time calibrated phylogenetic analyses in the future. Given that the order has been described by Silvestri in 1913 and 22 out of the 47 extant species have been described just in last 25 years (Choe, 2018; Kaláb et al., 2025; Kočárek & Horká, 2023a, 2023b; Lima et al., 2024; Matsumura et al., 2023), it is apparent that true evolutionary diversity of Zoraptera remains largely unknown (Choe, 2018).

### Chromosome number evolution

The estimated slopes of chromosome fission and fusion models and ancestral chromosome numbers support that karyotype evolution in Zoraptera is more frequently formed by chromosome fusions than fissions. The prevalence of chromosome fusions is evident in Zorotypinae subfamily where chromosome number decreased from Zorapteran ancestral 2n = ∼44 (MI) (or 2n = ∼52 as inferred by BI and MR) to 2n = 22 in *Zorotypus* sp. 1. It also aligns with the abundancy of ITSs in Spiralizorinae, since chromosome fusions are one of the main ITS origin mechanisms, as mentioned above. The estimated ancestral chromosome numbers (Fig. 9) had generally low statistical supports and rather serve as an approximate ancestral chromosome numbers. To refine the karyotype evolution history in Zoraptera, karyotype data of additional species and comparative genomic analyses are needed are needed in the future studies.

## Conclusions

In this study, we performed cytogenetic analysis of seven species of Zoraptera from both Spiralizoridae and Zorotypidae families. We reject the suggestion of holocentricity in Zoraptera, since it was not present in any of the analyzed species. Our results show that the striking morphological uniformity of its representatives is in sharp contrast with karyotype diversification. The structural karyotype differences between individual species are illustrated by various chromosome morphology and number as well as distribution of 18S rDNA and telomeric sequences. Our results confirm the previously suggested XY SCS in *U. hubbardi*, with 18SrDNA being localized in X chromosome. Also, we suggest the XY SCS is ancestral in Zoraptera. We hypothesize the SCS could be actually ancestral in Polyneoptera, though, this needs to be confirmed in future studies. The estimated chromosome number evolution model and relative abundance of ITSs suggest that karyotype evolution in Zoraptera is more frequently driven by chromosome fusions than fissions.

## Acknowledgements

Zoraptera material from Madagascar was collected as part of the research project “Étude à long terme de la biodiversité des groups choisis d’insectes: Coléoptères, Hétéroptères, Homoptères, Lépidoptères et quelques familles de Micro Lépidoptères nocturne dans les localités préalablement sélectionnées en considération de la recherche et la protection de la biodiversité dans les aires protégées de Madagascar. Analyse des risques potentiels d’influencer négativement la biodiversité dans les régions étudiées”, in cooperation with the Department of Entomology, University of Antananarivo (permit issued to M. Trýzna). Material from Brunei was collected with permission granted by the Ministry of Industry and Primary Resources, Brunei Darussalam (2013–2015; permit issued to P. Kočárek), and material from Panama with permission granted by the Ministry of Environment, Section on Access to Genetic and Biological Resources (SARGEB), Panama (2023; permit issued to P, Mückstein). We thank the following collaborators for collecting Zoraptera samples: Miloš Trýzna (Municipal Museum of Ústí nad Labem, Czech Republic) and Petr Mückstein (Nature Conservation Agency of the Czech Republic). We would like also to thank Mrs. Ony Rakotoarisoa (Director, Madagascar National Parks), Dr. Jean-Claude Rakotonirina, Dr. Victor Razafindranaivo, Prof. Lala Harivelo Ravaomanarivo Raveloson (University of Antananarivo) and Dr. Rodzay bin Haji Abdul Wahab (Universiti Brunei Darussalam) for supporting research projects.

