## Supplementary material for "Karyotype evolution of angel insects (Zoraptera)": Fig. S1

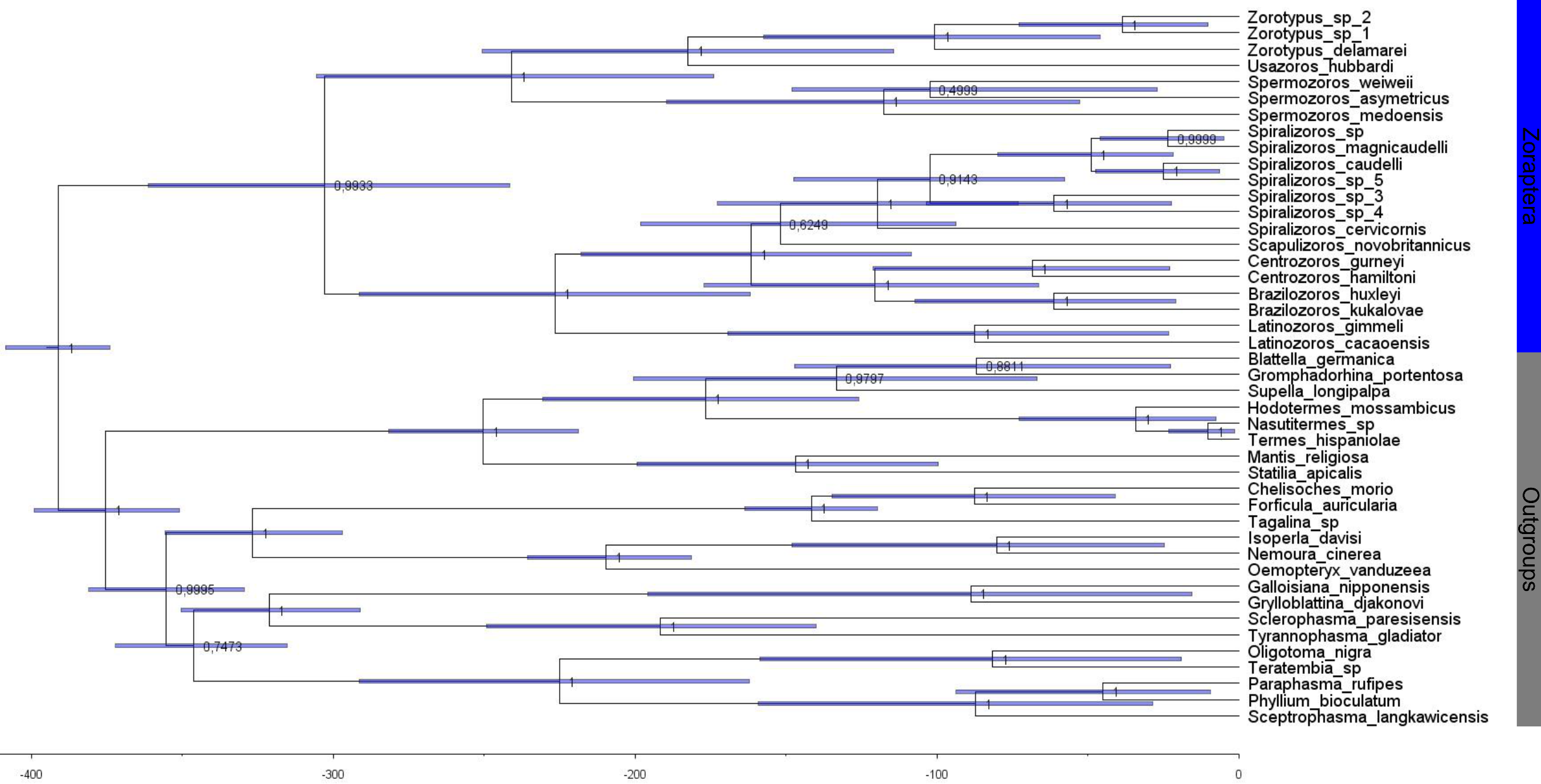

Figure S1: Time calibrated phylogenetic tree of Zoraptera produced by Bayesian inference method. Posterior probability is represented by the numbers next to the tree nodes. The blue bars around the tree nodes represent 95% confidence intervals.
